# ATG5 is dispensable for ATG8ylation of cellular proteins

**DOI:** 10.1101/2024.07.03.601942

**Authors:** Robin Ketteler, Koshiro Kiso, Lucas von Chamier, Alexander Agrotis

## Abstract

Protein ATG8ylation refers to a post-translational modification involving covalent attachment of ubiquitin-like autophagy-related protein ATG8 (LC3/GABARAP) to other cellular proteins, with reversal mediated by ATG4 proteases. While lipid ATG8ylation is important for autophagosome formation and mechanistically well-characterized, little is known about the mechanism of protein ATG8ylation. Here, we investigated the conjugation machinery of protein ATG8ylation in CRISPR/Cas9-engineered knockout human cell lines, utilizing a deconjugation-resistant (Q116P G120) form of MAP1LC3B. We report that protein ATG8ylation requires the E1-like activating enzyme ATG7 and E2-like conjugating enzyme ATG3, in common with ATG8 lipidation. However, in contrast, the E3-like ATG12-ATG5-ATG16L1 complex involved in lipidation is dispensable for protein ATG8ylation, since ATG5 knockout cells can form ATG8ylated protein conjugates. Further, we uncover that ATG7 itself is a target of ATG8ylation. Overall, our work provides crucial insight into the mechanism of protein ATG8ylation, distinguishing it from ATG8 lipidation, which will aid investigating its functional role.

## Introduction

Autophagy is an important intracellular process conserved in most eukaryotes, whereby damaged organelles, misfolded protein aggregates and/or other cytoplasmic cargoes are captured for lysosomal degradation [1,2]. The best-studied form of autophagy (macroautophagy, hereafter “autophagy”) utilizes autophagosomes, a type of double membrane organelle that is formed *de novo* to encapsulate material, through consecutive steps of protein kinase activation, lipid transfer and two ubiquitin-like conjugation reactions. Fully-formed autophagosomes eventually undergo SNARE-mediated membrane fusion with the lysosome to degrade their contents. Many of the proteins that coordinate autophagy are termed autophagy-related (ATG) proteins, which have been extensively studied in both mammalian and yeast cells.

Two ubiquitin-like conjugation reaction systems mediate autophagosome formation, comprising the ATG8 conjugation system and the ATG12-ATG5 conjugation system [3]. In the ATG8 conjugation system, ubiquitin-like ATG8 is covalently attached to phospholipids such as phosphatidylethanolamine (PE) on the forming autophagosome membrane. This process is termed lipidation or “membrane ATG8ylation”. In humans, ATG8 is encoded by at least seven gene orthologues including (MAP1)LC3A, LC3B, LC3B2, LC3C, GABARAP, GABARAPL1 and GABARAPL2, collectively referred to as “LC3/GABARAP” or simply “ATG8”. Directly assisting with membrane ATG8ylation in an E3-like manner, the ATG12-ATG5 conjugation system involves the covalent attachment of ubiquitin-like ATG12 to ATG5, which both then noncovalently associate with ATG16L1 to form the E3-like ATG16L1 complex (ATG12-ATG5-ATG16L1).

Like ubiquitylation, ATG8 lipidation is a multi-step process. ATG12 is conjugated to ATG5 through the sequential actions of the E1-like activating enzyme ATG7 and the E2-like conjugating enzyme ATG10. ATG8 also utilizes ATG7 as an E1-like enzyme, first binding to a region in the C-terminal domain of ATG7 in the two hydrophobic pockets of the ubiquitin-like fold [4]. ATG8 is transferred to the adenylation site of the ATG7 N-terminal domain containing the catalytic cysteine. ATG8 is next transferred to the active site of the E2-like conjugating enzyme ATG3, forming a thioester linkage. The final step of the ATG8 lipidation reaction, where ATG8 becomes attached to phospholipid headgroups, is catalyzed by the E3-like ATG12-ATG5-ATG16L1 complex [5]. ATG16L1 is the main determinant for targeting ATG8 to the membrane, aided by WIPI2 and PI3KC3-C1 [6,7].

A further layer of complexity exists through the reversibility of membrane ATG8ylation, mediated by the ATG4 cysteine proteases (ATG4A, ATG4B, ATG4C and ATG4D in humans) that cleave the C-terminus of ATG8. ATG8 is first synthesized with a pro-peptide that requires cleavage by ATG4 prior to conjugation, in a process known as ATG8 priming [8]. The fate of primed ATG8 can be quite diverse: in the canonical autophagy pathway it becomes attached to membrane during autophagosome formation, as described above [9,10]. ATG8 can also conjugate to single membranes of other organelles (termed “CASM”) [11], or even act as a transcriptional co-activator in the nucleus [12]. It is currently believed that priming is fast and constitutive [13] as opposed to being a regulatory step in ATG8ylation. However, since the activity of ATG4 proteases can be regulated by post-translational modifications [14–17], it is believed these might primarily serve to control the deconjugation (or delipidation) of ATG8.

Although the extent to which ATG4-mediated delipidation occurs during autophagy is unclear [13,18], in 2019, we discovered that ATG4 has an additional deconjugation role in removing ATG8 from proteins [19]. This was observed through expressing a deconjugation-resistant form of primed ATG8 (LC3B Q116P G120) in human cells, or expressing pre-primed ATG8 (LC3B G120 or equivalent truncations of each LC3/GABARAP) in cells lacking ATG4. In both cases, ATG8 was seen to attach to multiple cellular proteins which are substrates of ATG4, therefore establishing ATG8ylation (or “LC3ylation” as we first referred to it) as a novel reversible protein post-translational modification. Another research group published similar findings shortly after [20], and to date, protein targets of ATG8ylation reported include ATG3 [19], ATG16L1 [20] and NUFIP2 [21], although the functional role of this modification and full range of substrates remain enigmatic.

One important question is how protein ATG8ylation occurs, and which factors mediate or regulate this process. Here, we provide evidence that both ATG7 and ATG3 are required for the conjugation of ATG8 to proteins. However, in contrast to lipid conjugation, the ATG12-ATG5-ATG16L1 complex is not required. Therefore, we suggest that the targeting of membrane ATG8ylation and protein ATG8ylation are differentially controlled in cells, through the availability of the ATG16L1 complex. Finally, we establish ATG7 as a novel protein substrate of ATG8ylation.

## Results

ATG8ylation of membranes is mediated by ATG7, ATG3 and the ATG12-ATG5-ATG16L1 complex [3]. To determine whether the same components are required for protein ATG8ylation, we generated HeLa knockout (KO) cell lines lacking ATG5 or ATG7 protein using CRISPR/Cas9 genome editing, in addition to obtaining commercially-produced HAP1 *ATG5* KO cells, and using HeLa *ATG3* KO cells we validated in a previous study [19].

We confirmed loss of ATG5, ATG7 or ATG3 protein expression by western blotting in at least two HeLa knockout clones for each gene (Figure 1A). We also detected SQSTM1/p62 and LC3B to assess the effect of gene knockout on autophagy. As expected, lipidated LC3B-II could not be detected for all knockout clones of ATG3, ATG5 and ATG7, in contrast to wild-type HeLa cells, since conjugation of LC3B to membranes is defective in the absence of these ATG8 conjugation machinery components (Figure 1A). We also observed elevated basal levels of the autophagy cargo receptor SQSTM1 in most knockout clones, consistent with an impairment in basal autophagy due to ineffective ATG8 lipidation. As further expected, loss of *ATG5* led to an absence of the stable covalent ATG12-ATG5 protein conjugate, and the conjugation of ATG12 to ATG5 was defective in *ATG7* KO cells but not *ATG3* KO (Figure 1A). By assessing the effect of autophagy stimulation (using the potent mTOR inhibitor Torin1) and inhibition (using the late-stage blocker bafilomycin A1) in HAP1 *ATG5* knockout cells, it was clear that LC3B lipidation remained defective even under conditions where wild-type control cells exhibit high levels of LC3B lipidation (i.e. co-treatment with both agents, as seen in Figure 1B) Further, Torin1-induced degradation of SQSTM1 did not occur in *ATG5* KO HAP1 cells. Thus, all knockout cell lines behaved with the expected impairments in LC3B lipidation and autophagy.

**Figure 1:**
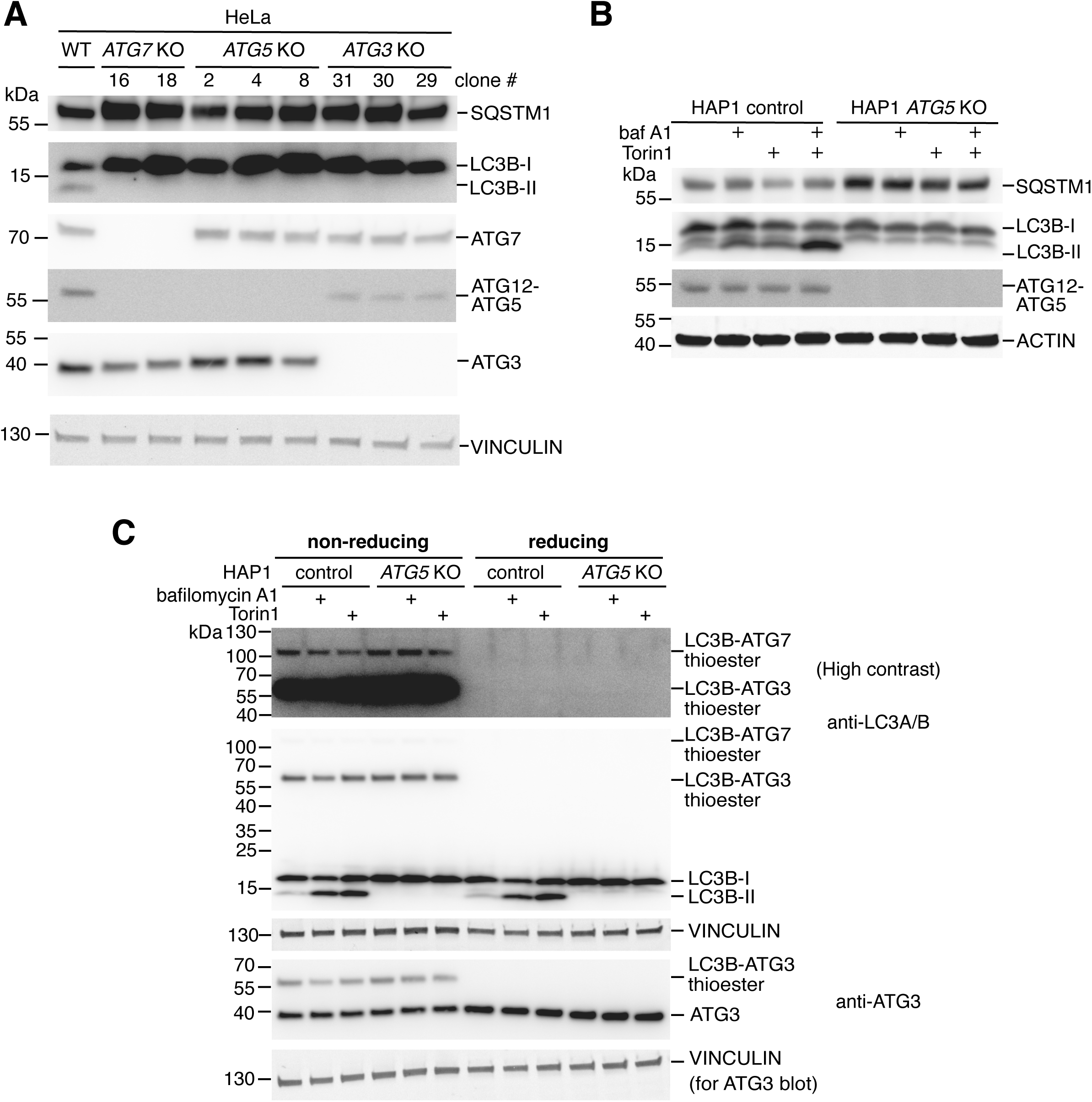
Characterization of human *ATG3*, *ATG5* and *ATG7* knockout cell lines. **A**, Western blot validation of CRISPR/Cas9-generated *ATG3*, *ATG5* and *ATG7* KO HeLa cell clones, using antibodies targeting ATG3, ATG5 and ATG7 protein. p62/SQSTM1 and LC3B were also detected to assess autophagy phenotypes. Vinculin was used as a loading control. **B**, Western blot analysis of the effect of autophagy inducer (Torin1, 250 nM) and inhibitor (baf A1; bafilomycin A1, 10 nM) single and combined treatments (for 3 hours prior to lysis) of HAP1 control and *ATG5* KO cells. Lysates were probed with antibodies against p62/SQSTM1 and LC3B, as well as ATG5 to detect ATG12-ATG5 conjugate. Actin was used as a loading control. **C**, HAP1 control and *ATG5* KO cells treated with Bafilomycin A1 (10 nM) or Torin1 (250 nM) for 3 hours were lysed, and samples were prepared for western blotting in the presence (reducing) or absence (non-reducing) of β-mercaptoethanol in the sample buffer. Membranes were probed with an antibody against ATG3 to assess total ATG3 expression and the redox-sensitive thioester linkage of LC3B with ATG3. An antibody against LC3A/B was used to visualize LC3B linked via thioester bond with ATG3 and ATG7 (visible only under non-reducing conditions) and LC3B lipidation. Vinculin was detected as a loading control.

We next tested whether *ATG5* knockout affects the ability of LC3B to bind to the active site of ATG7 or ATG3, which occurs through a redox-sensitive thioester linkage detectable by western blotting under non-reducing conditions [22] (Figure 1C). Indeed, despite being incapable of producing lipidated LC3B, HAP1 *ATG5* KO cells produced similar levels of LC3B-ATG3 and LC3B-ATG7 thioester compared to wild-type cells. Thus, loss of the ATG12-ATG5-ATG16L1 complex involved in ATG8 lipidation does not impact the transfer of LC3B to the upstream E1 and E2-like components of this pathway.

To investigate the mechanism of protein ATG8ylation, we made use of the LC3B Q116P G120 mutant that is resistant to deconjugation and is known to readily ATG8ylate other proteins when ectopically expressed in wild-type cells [19]. The pre-primed version of LC3B (G120) undergoes deconjugation by endogenous ATG4B in cells, and thus is less ATG8ylated, serving as a form of negative control in this experiment. We reasoned that a loss of deconjugation-resistant LC3B conjugates in a particular KO cell line would indicate that the missing gene is important for ATG8ylation. When expressed in HAP1 *ATG5* KO cells, LC3B Q116P G120 was capable of forming high molecular-weight protein conjugates at levels similar to HAP1 control cells (Figure 2A), suggesting that ATG5 is not required for protein ATG8ylation. We also observed the same phenotype in HeLa cells, since HeLa *ATG5* KO cells were clearly capable of forming LC3B protein conjugates (Figure 2B). In contrast, there was a marked deficiency in LC3B Q116P G120 conjugate formation in HeLa *ATG3* and *ATG7* KO cells (Figure 2B). Altogether this implicates ATG7 and ATG3, but not ATG5, in the process of ATG8ylation.

**Figure 2:**
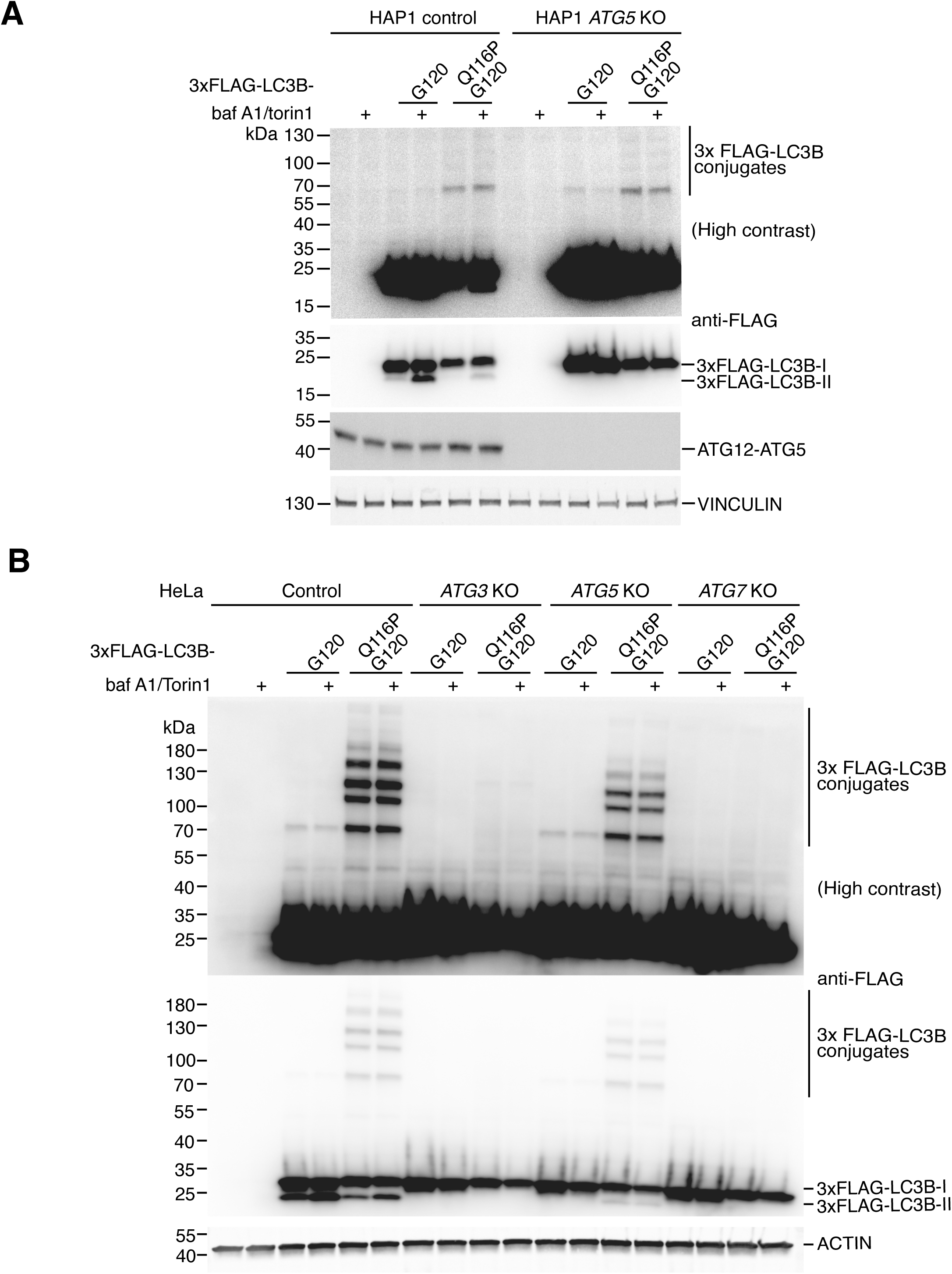
ATG5 is dispensable for protein ATG8ylation. **A**, HAP1 control and *ATG5* KO cells transfected with 3xFLAG-LC3B G120 or 3xFLAG-LC3B Q116P G120 were treated with DMSO or 250 nM Torin1 and 10 nM bafilomycin A1 for 3 hours prior to lysis. Lysates were analyzed by western blotting using antibodies against the FLAG tag, ATG5 (to detect ATG12-ATG5 conjugate) or Vinculin (as a loading control). **B**, HeLa control, *ATG3* KO, *ATG5* KO or *ATG7* KO cells were transfected with Flag-tagged LC3B G120 or LC3B Q116P G120 and treated for 3 hours with DMSO or or 250 nM Torin1 and 10 nM bafilomycin A1 prior to lysis. western blotting of lysates was performed and the membrane was probed with antibodies against FLAG tag, or Actin as a loading control.

When further studying the functional link between ATG7 and ATG8ylation, we noted a specific accumulation of two additional high molecular-weight bands (between 100-130 kDa) when blotting for ATG7 in ATG4B knockout cells expressing LC3B G120 i.e. under conditions where ATG8ylated proteins are known to accumulate [19] (Figure 3A). To validate whether these bands represented ATG8ylated ATG7, we purified LC3B conjugates from ATG4 KO cells expressing LC3B G120 and Q116P G120 by immunoprecipitation and blotted for ATG7 following digestion with recombinant ATG4B (or catalytic inactive ATG4B C74S as a negative control). Our evidence suggests that these two bands indeed correspond to LC3B conjugated to ATG7 in the form of ATG8ylation, since the LC3B G120 forms of these bands could be abolished specifically by treatment with active ATG4B, but the bands were resistant to deconjugation when LC3B Q116P G120 was used (Figure 3B). We also detected the known ATG8ylation substrate ATG3 [19] in the same samples as an internal validation for our IP-digestion assay, which behaved as predicted, with LC3B G120 conjugates being cleaved by active recombinant ATG4B (Figure 3B). Furthermore, since western blotting was performed under reducing conditions, this confirms that ATG8ylated ATG7 is distinct from the ATG8-ATG7 thioester which forms as a part of the ATG8 conjugation mechanism.

**Figure 3:**
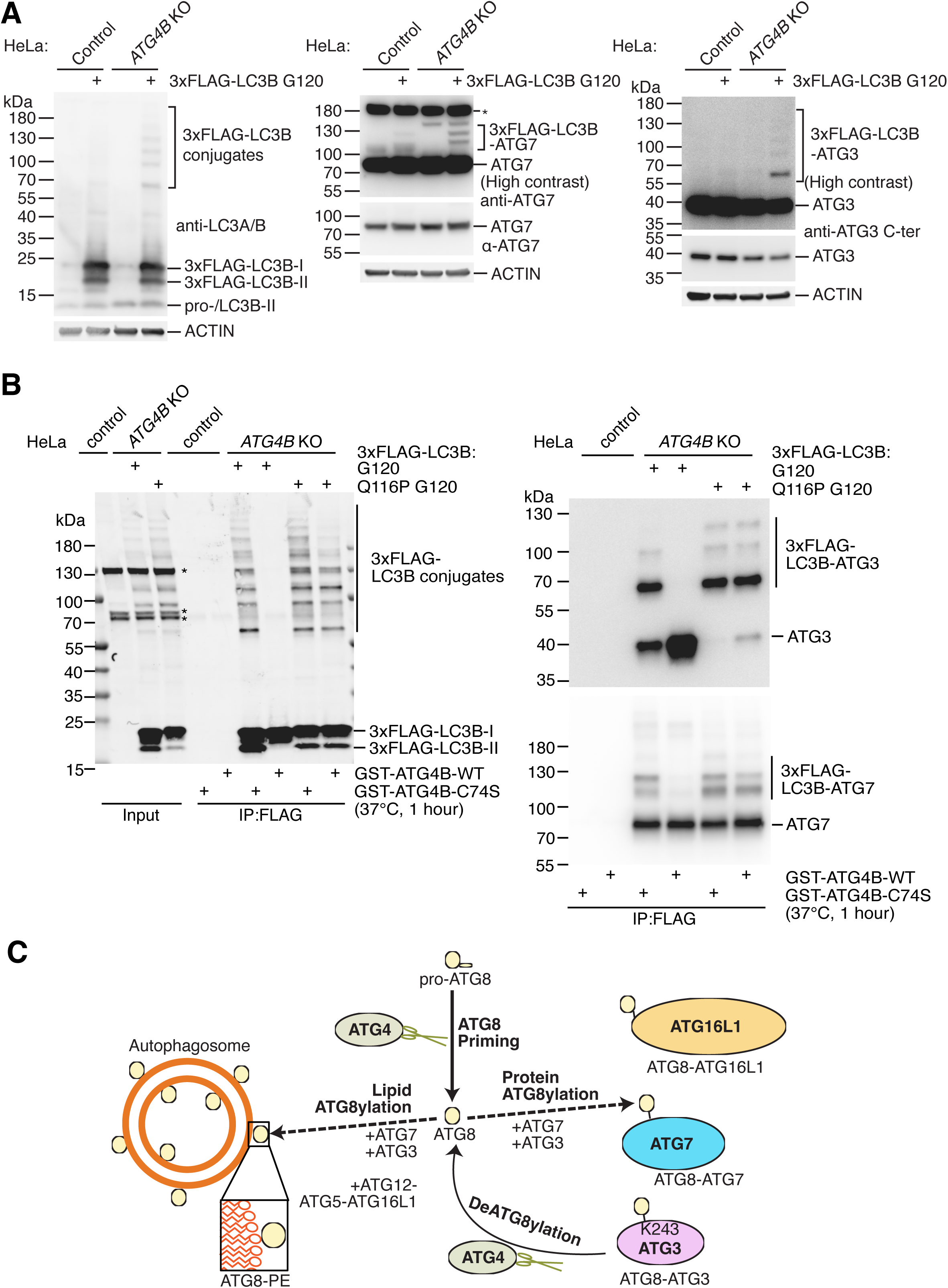
ATG7 is a protein target of ATG8ylation. **A**, HeLa control and *ATG4B* KO cells were transfected with 3xFLAG-LC3B G120 (or mock transfected as a negative control) and lysates were subject to western blotting with anti-LC3A/B, anti-ATG7 and anti-ATG3 C-ter antibodies, with Actin detected as a loading control. Blots for ATG3 and ATG7 are additionally presented as high contrast images to improve visibility of less prominent bands (*asterisk denotes non-specific band). **B**, HeLa control cells were transfected with GFP as a negative control and HeLa *ATG4B* KO cells were transfected with either 3xFLAG-LC3B G120 or 3xFLAG-LC3B Q116P G120. Cells were lysed in IP buffer and subject to immunoprecipitation with anti-FLAG® M2 Affinity Gel. Eluates were digested with either purified recombinant GST-ATG4B WT or GST-ATG4B C74S protein added to reactions at a final concentration of 0.02 mg/ml and incubated for 1 hour at 37°C prior to inactivation with sample buffer and boiling at 95°C for 5 minutes. Samples including input and IP samples were analyzed by western blotting using anti-FLAG antibody (*non-specific bands are indicated with asterisks) and immunoprecipitated samples on separate membranes were probed with anti-ATG3 and anti-ATG7 antibodies. **C**, Summary of updated model of ATG8ylation incorporating key findings from this study. There are distinct mechanisms of lipid ATG8ylation (requiring the ATG12-ATG5-ATG16L1 complex) versus protein ATG8ylation (which is independent of ATG5). ATG4 proteases carry out deATG8ylation of proteins including ATG3, ATG16L1 and ATG7.

Overall, our data indicate that ATG8ylation of proteins requires ATG3 and ATG7, both of which are also targets for ATG8ylation. In contrast, the presence of the ATG12-ATG5-ATG16L1 complex is not required for protein ATG8ylation, providing a distinction between the mechanism of lipid versus protein ATG8ylation (Figure 3C).

## Discussion

Membrane ATG8ylation is important in many processes linked to human health and disease, including LAP (LC3-associated phagocytosis), starvation-induced autophagy and selective forms of autophagy. It has been proposed that ATG8ylation in general may be a marker of cellular stress, with varying functions in processes such as energy homeostasis, starvation, immune defense and other forms of stress [23–26].

Since the discovery that proteins can be ATG8ylated and subsequent studies confirming additional targets of protein ATG8ylation [19,20], little has been reported about the machinery involved in ATG8ylation of proteins. We demonstrate here that the same E1- and E2-like enzymes that are involved in lipid ATG8ylation are also required for protein ATG8ylation. However, in contrast to membrane ATG8ylation, the ATG12-ATG5-ATG16L1 complex is not required for protein ATG8ylation, suggesting distinct regulatory mechanisms. There are several implications following from this finding, for example, it may be the case that the actions of ATG7 and ATG3 alone are sufficient for protein ATG8ylation. *In vitro*, it has been shown that ATG7 and ATG3 are sufficient for the transfer of ATG8 to PE on small unilamellar vesicles containing a high percentage of PE[3,27]. The addition of ATG12-ATG5-ATG16L1 greatly enhances this lipidation *in vitro* and is required for lipidation in cells [4,5,28–31]. It is conceivable that ATG7 and ATG3 are sufficient for transfer of ATG8 to targets in general, while in cells there may be additional factors required to facilitate such transfer to lipids. Considering that the transfer of ATG8 to membranes is highly dependent on the curvature and composition of the membrane [32], it may be that ATG8ylation of intracellular proteins has fewer constraints, and thus ATG3 and ATG7 could suffice. However, our results do not rule out the potential presence of other E3-like components required protein ATG8ylation, therefore further experiments, for instance genetic screening or proteomic approaches, could be utilized to explore this possibility.

In the ubiquitin-proteasome system, it is generally thought that the E3 ligase determines substrate specificity, although in some cases protein ubiquitylation can occur without an E3 ligase [33,34]. It is possible that by analogy, for ATG8ylation, substrate specificity is determined by the E3-like complex. As mentioned, proper membrane ATG8ylation is mediated by the activity of the ATG16L1 complex, while substrate-agnostic default ATG8ylation of other targets (i.e. proteins) may be mediated by ATG7 and ATG3 alone. Supporting this notion are recent observations that the composition of the ATG16L1 complex mediates substrate specificity. For instance, membrane binding of ATG5 is required for PE lipidation [5] and enhanced by the presence of WIPI2 [6,7]. In general, in the presence of ATG16L1 protein, ATG8ylation is directed towards autophagosomal membranes, whereas in an ATG12-ATG5 complex containing TECPR1 instead of ATG16L1, ATG8ylation is directed towards sphingosine-containing membranes [35,36]. In addition, the WD40 repeat region in ATG16L1 directs ATG8ylation towards single membranes during LAP [31,37]. Thus, one could hypothesize that co-factors in the ATG12-ATG5 E3-like complex have a key role in determining substrate specificity [38].

What is the function – if any – of protein ATG8ylation? Several hypotheses have been brought forward, including a function in stress response, or serving as a reservoir for ATG8 that can be quickly mobilized. To date, only ATG3 [19], ATG7 (this study), ATG16L1 [20], and NUFIP2 [21] have been identified as protein ATG8ylation targets. All except NUFIP2 are key regulators of the ATG8ylation process itself, raising the possibility that ATG8ylation is a by-product that occurs during the ATG8ylation of membranes. It should be noted that the protein ATG8ylation we refer to in this study is distinct from the thioester linkage between ATG8 and the catalytic site of ATG7 or ATG3. In yeast, it has been observed that ATG4 proteases act to remove aberrantly targeted ATG8 [39], thus the simplest explanation might be that proteins are just aberrantly targeted by ATG8 when autophagosomes are not present in sufficient numbers or when the ATG12-ATG5-ATG16L1 complex is missing. Perhaps “aberrant” targeting of proteins by ATG8ylation serves as a reservoir whereby ATG8 can be quickly mobilized by ATG4 proteases. Interestingly, it has been suggested that non-lipidated GABARAP at the centrosome serve as a reservoir of ATG8 [40]. Alternatively, ATG8ylation of ATG3 and ATG7 might serve as a feedback mechanism to regulate their activity, and this could be the subject of future research.

Whether additional proteins exist that are ATG8ylated remains to be determined. We propose that the expression of deconjugation-resistant ATG8 and pre-primed ATG8 in ATG4 deficient cells, combined with digestion using recombinant ATG4 following substrate enrichment by immunoprecipitation (as demonstrated in Figure 3B), should be considered the gold standard method for validating novel ATG8ylated substrates. The identification of the ATG8ylated proteome will provide further clues about the potential function and relevance of this novel post-translational modification of proteins in cells.

## Material and Methods

### Cell culture

Cells were cultured at 37°C, 5% CO^2^. HeLa cells were grown in DMEM high glucose with GlutaMax and 1 mM pyruvate (ThermoFisher Scientific, 31966021), supplemented with penicillin-streptomycin (ThermoFisher Scientific, 15140122; 100 U/ml each) and 10% FBS (ThermoFisher Scientific, 10270106). HAP1 cells were grown in base medium of IMDM containing 25 mM HEPES and L-glutamine (ThermoFisher Scientific, 12440053) supplemented as above for HeLa cells.

CRISPR/Cas9-mediated knockout of *ATG7* and *ATG5* in HeLa cells was performed using the method previously described for HeLa *ATG3* KO cells [19]. The oligonucleotides used to target exon 4 of *ATG7* (location: 3:11306996-11307018) were: sgATG7(+) (5’-CACCGTTGAAAGACTCGAGTGTGT-3’) and sgATG7(-) (5’-AAACACACACTCGAGTCTTTCAAC-3’), and oligonucleotides targeting exon 5 (location: 6:106279672-106279694) of *ATG5* were: sgATG5(+) (5’-CACCGATCACAAGCAACTCTGGAT-3’) and sgATG5(-) (5’-AAACATCCAGAGTTGCTTGTGATC-3’). These oligonucleotide pairs were annealed, phosphorylated, and cloned into BbsI-digested pSpCas9(BB)-2A-Puro (PX459) version 2.0, a gift from Feng Zhang (Addgene plasmid no. 62988) followed by transient transfection in wild-type HeLa cells using JetPRIME. At one day post-transfection, cells were grown for 48 h in the presence of 1 μg/ml puromycin. Surviving cells were then expanded into 10 cm dishes for 72 h, prior to seeding at a limiting dilution in 96-well plates. After 2 weeks, clonal cell lines were expanded into 12-well plates and screened for loss of ATG7 and ATG5 protein by Western blotting. Unless otherwise stated, the main knockout clones used for experiments were HeLa *ATG5* KO (c4), *ATG7* KO (c16) and *ATG3* KO (c31, as generated previously [19]). HeLa *ATG4B* KO cells were generated and characterized by our lab in a previous study [18]. HAP1 control and *ATG5* KO cells (cat no: HZGHC016335c010) were obtained from Horizon Genomics.

HAP1 cells were transfected with Viromer Yellow (Lipocalyx GmbH) according to the manufacturer’s protocol using 1 µg total DNA per 12-well plate well containing 1 ml complete growth medium, and medium was replaced at 4 hours post-transfection. HeLa cells were transfected with 0.25 µg of DNA and 1.5 µl of jetPRIME reagent (Polyplus transfection) mixed in 50 µl jetPRIME buffer per 12-well plate well containing 1 ml growth medium. This was scaled up according to growth medium volume of the vessel used. All experiments were performed 24 h after transfection.

### Western blotting

Western blotting was performed exactly as described in detail previously[19]. In brief, cells were washed in PBS and lysed in lysis buffer (150 mm NaCl, 1% IGEPAL CA-630, 50 mm Tris-HCl, pH 8.0, cOmplete EDTA-free Protease Inhibitor Mixture, Roche Applied Science) supplemented with 20 mm N-ethylmaleimide (NEM), except experiments comparing reduced vs non-reduced samples where NEM was excluded. Lysates were cleared by centrifugation and diluted to equal total protein concentrations following analysis using Pierce BCA protein assay kit (Thermo Fisher Scientific). Prior to loading on gels, samples were mixed with sample buffer (to a final concentration of 50 mm Tris-Cl, pH 6.8, 2% SDS, 10% glycerol, 5% β-mercaptoethanol, and 0.01% bromphenol blue) and boiled at 95°C for 5 minutes. Samples of 15-30 µg total protein were run on 15-well 4–20% Mini-PROTEAN TGX Precast gels or 26-well 4–20% Criterion TGX gels using the Tris-Glycine buffer system and compatible running and transfer tanks for wet transfer (Bio-Rad). Samples were transferred to Immobilon-FL PVDF membrane (Merck-Millipore) prior to blocking in 5% milk in PBS-T (0.1% Tween-20), overnight primary antibody incubation at 4°C, washing in PBS-T and secondary antibody incubation for 1 hour at room temperature. After washing, blots incubated with HPR-conjugated secondary antibodies were developed using HRP substrate (EZ-ECL chemiluminescence detection kit for HRP, Geneflow Ltd.) followed by exposure on an ImageQuant chemiluminescent imaging system (GE Healthcare). For blots probed with fluorescent secondary antibodies, detection was performed using an Odyssey IR imaging system (LI-COR Biosciences).

### Immunoprecipitation

Immunoprecipitation was performed exactly as described in detail previously [19]. In short, cells grown and transfected in 10 cm dishes were washed in PBS and lysed in IP buffer (50 mm Tris-HCl, pH 8, 150 mm NaCl, 1 mm EDTA, 1% Triton X-100, and cOmplete™ EDTA-free Protease Inhibitor Mixture, Roche Applied Science) before clarification by centrifugation and enrichment for FLAG-tagged proteins using anti-FLAG® M2 Affinity Gel (A2220, Sigma), with elution using 3xFLAG peptide (F4799, Sigma) at 150 ng/µl in IP buffer. Immunoprecipitated samples were subjected to treatment with recombinant GST-ATG4B or catalytic inactive GST-ATG4B C74S, mixed with sample buffer and boiled, and analyzed by western blotting.

### Antibodies, plasmids and reagents

Primary antibodies and dilutions used for western blotting in this study were: ACTIN (1:4,000, Sigma, A1978), ATG3 (1:1,000, Abcam, ab108251), ATG3 C-ter (1:1,000, Abcam, ab108282), ATG5 (used to detect ATG12-ATG5 conjugate; 1:1000, Cell Signaling Technology, no. 2630), ATG7 (1:2,000, Abcam, ab52472), FLAG biotin-conjugated (1:1,000, Sigma, F9291), LC3A/B (1:500, Cell Signaling Technology, no. 12741), SQSTM1 (1:1,000, Sigma, P0057) and VINCULIN (1:1,000, Abcam, ab129002).

Secondary antibodies used were: goat anti-rabbit/anti-mouse IgG, HRP-linked (1:5,000, Cell Signaling Technology, no. 7074/no. 7076); IRDye 800CW goat anti-rabbit/anti-mouse (1:15,000 with 0.02% SDS, LI-COR no. 926-32211/926-32212); IRDye 680LT goat anti-rabbit/anti-mouse (1:25,000 with 0.02% SDS, LI-COR no. 926-68021/926-68020); IRDye 680LT Streptavidin (1/5,000 with 0.02% SDS, LI-COR 926-68031).

Plasmids used in this study were generated by our lab previously: pCMV 3xFLAG-LC3B G120 (Addgene plasmid # 123094; http://n2t.net/addgene:123094) [18] and pCMV 3xFLAG-LC3B Q116P G120 (Addgene plasmid # 129289; http://n2t.net/addgene:129289) [19]. pEAK13-EGFP was used for expression of GFP alone [19].

Torin1 from Merck-Millipore (no. 475991), and *Streptomyces griseus* bafilomycin A1 from Sigma (B1793) were dissolved in DMSO. DMSO was used as a negative control for treatment experiments. Recombinant GST-ATG4B WT and C74S were produced in a previous study [41].

## Acknowledgements

This work was supported by UK Medical Research Council core funding to the MRC-UCL University Unit Grant Ref MC_U12266B, MRC Dementia Platform Grant UK MR/M02492X/1, the UCL Therapeutic Acceleration Support scheme, supported by funding from MRC Confidence in Concept 2020 UCL MC/PC/19054 and Wellcome Trust Grant 105604/Z/14/Z. This article/publication is based upon work from COST Action ProteoCure, CA20113, supported by COST (European Cooperation in Science and Technology). We thank Oscar Blackwell for manuscript feedback.

